# Developmental Changes in Gray Matter Microstructure and Local Brain Connectivity Support Inhibitory Control from Adolescence to Young Adulthood

**DOI:** 10.64898/2026.08.19.745824

**Authors:** Victoria O. Dionisos, Valerie J. Sydnor, Will Foran, Finnegan J. Calabro, Beatriz Luna

**Author notes:** Correspondence: Victoria O. Dionisos, Laboratory of Neurocognitive Development University of Pittsburgh, 121 Meyran Ave, Pittsburgh, PA 15213.

## Abstract

**Background:** Adolescence is marked by improvements in inhibitory control along with brain maturational processes affecting function and structure of cortical circuitry. Preliminary evidence shows a developmental decrease of local neuronal inputs, suggesting a weakening of local connectivity supporting circuit refinement and mature behavior. Microstructural changes in gray matter, arising from processes such as synaptic pruning and myelination, may support this refinement, though it remains unknown how microstructural features interact with the reconfiguration of functional circuitry, or how this unfolds *in vivo* in normative development to support mature cognitive functioning.

**Methods:** In this study, 175 participants ages 10-26 (93F; 17.32±4.82yo) completed an anti-saccade task, a developmentally-validated measure of inhibitory control, as well as a 3T MRI scan involving a multi-shell diffusion weighted imaging acquisition, multi-echo resting state fMRI, and structural (T1w, T2w) scans. We computed measures of neurite density (NDI) in gray matter using neurite orientation dispersion and density imaging, local functional connectivity with surface-based and volumetric regional homogeneity (ReHo), and intracortical myelin using T1w/T2w ratio. Generalized additive models examined non-linear age-related trends across a number of cortical and subcortical regions implicated in inhibitory control, as well as associations with anti-saccade performance.

**Results:** We found that NDI significantly increased with age in all regions while ReHo decreased. Greater NDI was associated with more accurate anti-saccade performance and lower ReHo, which remained after residualizing NDI for T1w/T2w ratio, suggesting that microstructural reorganization beyond myelination may underlie functional specialization throughout adolescence. Lower ReHo, specifically in young adolescents, also resulted in better anti-saccade performance. Finally, an interaction between ReHo and NDI, rather than either measure alone, best predicted inhibitory control performance, such that the maturity of neurite density had the greatest effect when local connectivity was high, suggesting immaturity.

**Conclusions:** Our results suggest that the joint maturation of microstructural elements and associated specialization of local functional circuitry interact to support the emergence of stable adult-level inhibitory control.

## Introduction

The adolescent period is marked by improvements in complex information processing and executive functioning. Inhibitory control, a core process of executive function, requires the coordination of cognitive and motor systems to effectively inhibit prepotent responses in favor of an executive, goal-directed response ^1^. Previous studies with non-human primates and humans have shown that inhibitory control improves with age ^1–7^, which continues through the second decade of life before beginning to plateau and stabilize in adulthood ^3,5^. This ability to stably and flexibly control behavior is supported by refinements in activation and communication across executive and motor control regions. This includes prefrontal and parietal regions, including the dorsal anterior cingulate cortex, dorsolateral prefrontal cortex, frontal/supplementary eye fields, posterior parietal cortex and pre-supplementary motor area, along with subcortical regions including the caudate and putamen ^5,8^. Many of these regions communicate with each other via a cognitive and motor loop through the basal ganglia, with frontal oculomotor regions connecting directly to the caudate, and sensorimotor regions connecting to the putamen ^9^. All together, these regions play a crucial role in coordinating goal-directed responses and guiding top-down processing necessary for mature cognitive control ^10^, which is integral for flexible updating and planning of behavior recruited during inhibitory control ^5^.

In parallel with developmental improvements in executive function are brain maturational processes that affect structure and function of cognitive circuitry. Protracted pruning of excitatory synapses in the prefrontal cortex begins around the start of puberty in early adolescence and continues into adulthood ^11–13^. This selective elimination of redundant or inefficient synapses has been proposed as a core developmental process guiding the reorganization of cortical and subcortical circuitry ^13^. Significant evidence in animal and post-mortem human tissue studies have shown that synaptic pruning directly shapes the underlying microstructure surrounding pruned synapses, allowing for optimization of information processing to directly support complex processes such as cognitive control ^14^. Across development, synaptic refinement results in a reduction of dendritic spines and dendritic arborization ^13,15^, as well as increased myelin ^16,17^. However, due to methodological limitations, these microstructural changes have yet to be comprehensively characterized *in vivo* in human adolescent development.

Modeling approaches based on diffusion weighted imaging (DWI), such as neurite orientation dispersion and density imaging (NODDI), can be applied to assess these tissue properties *in vivo*. NODDI measures the diffusion rate of water across tissue compartments, producing an intra-neurite volume fraction, or neurite density index (NDI), which reflects the density of myelinated and unmyelinated axons and dendrites within a specific voxel ^18,19^. This provides an index of how tightly packed neuronal elements are in a given space. Preliminary evidence has shown that NDI increases during development in both cortical and subcortical gray matter ^20–23^ and is highly sensitive to age compared to traditional DWI measures ^24–27^. NDI is thought to reflect a combination of structural features, including a strong contribution from intracortical myelination, as it has been shown to highly correlate with myelin density estimated from T1w/T2w images ^28^ and histological myelin ^29^. Given that myelin increases through adolescence in a hierarchical manner with frontal regions continuing to myelinate into young adulthood ^17,30–33^, developmental increases in myelin may mask other microstructural properties characterized by NDI, including axonal growth and changes in axonal packing, dendritic branching, or density of microglia associated with pruning.

Alongside changes in microstructure are changes in function, which are known to be highly interdependent on each other ^15,34^. Micro-level structural changes shaped by synaptic plasticity, including pruning, have been linked to changes in micro-level or regional functional properties. For example, evidence in rodents demonstrates a selective developmental reduction in local excitatory synapses, which innervate dendrites proximal to the neuronal body and connect intra-regionally, leading to more optimal cross-region information processing supporting mature cognitive behavior via sparse local connectivity ^35,36^. Recent human neuroimaging studies mirror this, demonstrating developmental reductions in regional homogeneity, a neuroimaging measure of local, intra-regional functional connectivity assessing how correlated a single point is to its closest neighbors ^37,38^. These cross-species findings reflect a trend toward region specialization during development as local connections become sparser and more functionally independent ^37^. Given that changes in microstructure and local functional connectivity occur in parallel and may interact to guide development, we sought to understand the maturation of both structure and function at the regional level supporting developmental improvements in inhibitory control.

The current study investigated how micro-level structural and functional organization jointly support normative development of inhibitory control. To do so, we first aimed to characterize developmental trajectories of gray matter microstructure and local functional connectivity in oculomotor networks known to support inhibitory control, in addition to how each respectively relates to improved cognitive performance. Based on preliminary evidence, we predicted that neurite density would increase with age ^21–23^ whereas regional homogeneity would decrease ^37,39^; moreover, we hypothesized that both measures would be associated with improvements in cognitive functioning ^37,40^. Next, we evaluated the role of microstructural changes in local functional connectivity changes, and how they interact to influence inhibitory control. Since neurite density and regional homogeneity are both measured at the micro (intra-regional) level, we expected significant interactions among these measures, including in their contribution to the maturation of inhibitory control.

## Methods and Materials

### Participants

175 participants ages 10-26 (93F; 17.32±4.82yo) were recruited from the greater Pittsburgh area. Exclusion criteria were as follows: participant diagnosis or family history of psychiatric or neurological disorder, current use of psychiatric medication, history of head injury with loss of consciousness, vision problems, history of drug abuse, learning disability, or MRI counterindications. Demographic distributions of race and ethnicity included: 11% Asian, 4.5% Black, 8% Multiracial, 75% White, and < 2% did not specify, with 95.5% self-reporting as non-Hispanic. Date of birth and other demographic data was collected at the point of enrollment. Study procedures were approved by the University of Pittsburgh institutional review board and carried out in accordance with the Declaration of Helsinki. Adult participants gave written, informed consent to enroll in the study; participants under 18 could not enroll without written assent and a legal guardian’s written consent. Participants were compensated for their completion of each study visit.

### Procedures

#### Behavioral Data Acquisition

Participants completed an electro-oculogram (EOG)-monitored anti-saccade task, a well-studied measure of inhibitory control ^41^, during an electroencephalography session (EEG) (See Figure 1A). Participants were seated in front of a computer screen and instructed that when a visual stimulus appears in their periphery, they should suppress the prepotent response to look at the stimulus and instead direct their gaze to the mirror location of the stimulus ^1,6^. Trials began with an inter-trial interval ranging from 500-6000ms with a white fixation cross, followed by a preparatory epoch where a red fixation cross appeared for 200ms. At the extinction of the red cross, a yellow circular cue was presented in an unpredicted location along the horizontal axis (± 8°, 16° or 23° visual angle relative to fixation) for 1000ms, during which the participant must generate a saccade to the mirror location. Successful execution of the task demands both the suppression of a prepotent response and the generation of a voluntary, goal-directed oculomotor response^1^. An error on the task consists of making a saccade towards the stimulus then quickly correcting with a second saccade to the mirror location of the stimulus (“error corrected”), indicating the participant’s understanding of the task but their inability to inhibit their reflexive response and engage a goal-directed response. Two 10-minute runs of the task were administered with EOG recording for tracking eye movements from electrodes placed on the outer edge of each eye. A 40-position calibration occurred before each session to generate a participant-specific slope relating EOG signal to screen position.

**Figure 1.**
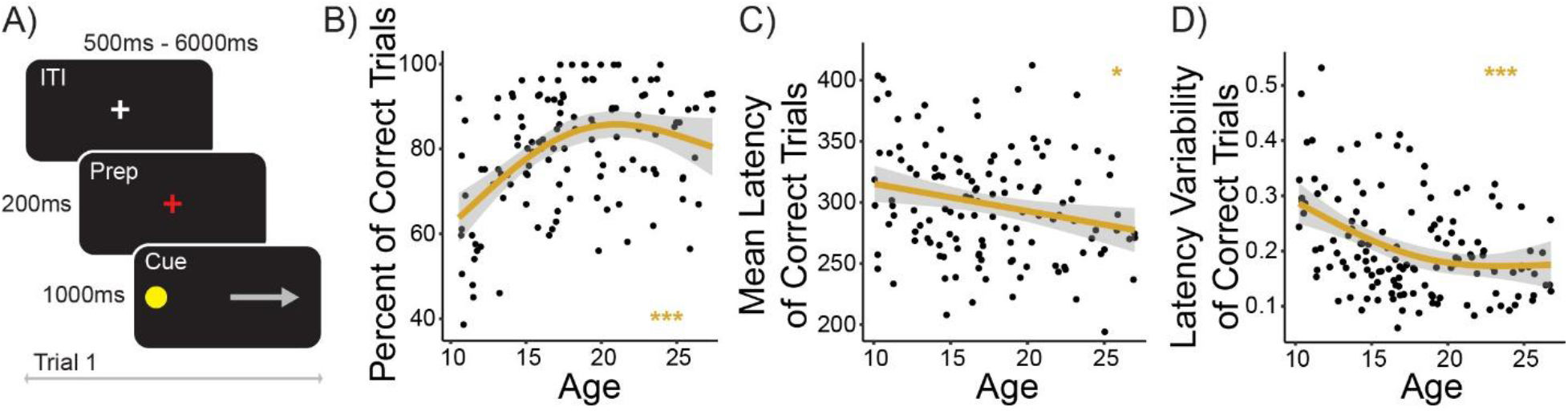
Developmental improvements in the anti-saccade task. **A)** Anti-saccade task design. Inhibitory control improved with age, reflected by improvements in the **B)** proportion of correct inhibitory responses, **C)** response time (latency) on correct trials, and **D)** trial-to-trial variability (standard deviation) of response time on correct trials. Significance survived FDR correction for multiple comparisons. *\*\*\*p* < .001, *\*p* < .05.

#### Behavioral Data Processing

EOG recordings from the anti-saccade task were evaluated with per-session calibrations via MATLAB to calculate saccade onset and velocity. Saccade information was used to score outcomes per trial, each 1 second in duration. All trials were scored as either being dropped (excluded), incorrect, correct, or error corrected. Trials were excluded if the first saccade occurred earlier than 100 milliseconds or if the saccade velocity before the stimulus appeared was greater than a set threshold (30°/sec), as this reflects an express saccade that did not engage executive systems ^42^. Percent of correct trials, calculated out of correct and error-corrected trials and reflecting when participants successfully inhibited a response, as well as mean latency and latency variability of correct trials were calculated for analysis. Analyses excluded participants that had fewer than 14 viable trials (out of 28) and less than 50% “on-task” trials, which excluded dropped and incorrect trials. Following these exclusions, the final sample was n=134 (10-26yo; 70F).

#### MR Data Acquisition

At a separate visit, participants completed a 3T neuroimaging scan (SIEMENS MAGNETOM Prisma_fit; Siemens Medical Solutions; Erlangen, Germany) using a 32-channel head coil involving multi-shell diffusion weighted imaging (DWI), multi-echo resting state (rs-fMRI), and structural (T1w, T2w) scans at the UPMC Magnetic Resonance Research Center. T1w and T2w image acquisition parameters from the ABCD Study were utilized ^43^. The T1w image was used for alignment to standard MNI space (MNI2009c) with the following parameters: TR = 2500ms, flip angle = 8°, voxel size 1×1×1mm, TI = 1070ms, TE = 2.9ms, 1mm^3^ isotropic resolution, GRAPPA acceleration factor = 2, scan time = 7 min 12 sec. The T2w scan was acquired with the following parameters: TR = 3200ms, voxel size 1×1×1mm, TE = 565ms, 1mm^3^ isotropic resolution, GRAPPA acceleration factor = 2, scan time = 6 min 35 sec. Multi-shell DWI (b=1500/3000 s/mm^2^, dir=93/92) with 12 non-diffusion weighted images (b = 0 s/mm^2^) had the following parameters: TR = 3230ms, TE = 89.20ms, voxel size 1.5×1.5×1.5mm, total scan time = 11 min 07 sec. One non-diffusion sensitized spin echo volume was acquired for susceptibility distortion correction in both anterior to posterior and posterior to anterior directions; TR = 3230ms, TE = 89.20ms, voxel size 1.5×1.5×1.5mm, total scan time = 1 min 13 sec. Two multi-echo resting state acquisitions were completed with the following parameters: TR=1200ms, flip angle = 60°, voxel size = 3×3×3mm, TE = 14, 31.63, 49.26ms, GRAPPA acceleration factor = 2, multi-band acceleration factor = 3, scan time = 7 min 54 sec. All MR data were mapped to a gray matter-specific ROI mask in MNI2009c that included cortical and subcortical regions (see details below).

#### DWI Data Processing

We derived neurite density index (NDI) from DWI. B0 field map files for each run were matched with corresponding Anterior-Posterior and Posterior-Anterior runs. Data were processed through QSIPrep (version 1.0.0) and QSIRecon pipelines ^19,44^. QSIPrep processing steps included distortion grouping, denoising, tissue segmentation, head motion correction, susceptibility correction, co-registration to T1w images, and spatial normalization. The output of QSIPrep was visually inspected before continuing to QSIRecon post-processing. The QSIRecon docker script [https://qsirecon.readthedocs.io/en/latest/; github.com/PennLINC/qsirecon] was executed with the command-amico_noddi to compute neurite density using the NODDI model and AMICO algorithm ^45^. Again, we visually inspected the QSIRecon output, followed by warping our ROI mask in MNI to ACPC subject space using antsApplyTransforms and extracting ROIs with AFNI’s 3dROIstats. All scans with framewise displacement greater than 1 were excluded from analysis, making the final sample n=100 (10-26yo; 55F).

#### rs-fMRI Data Processing

Local connectivity was characterized by regional homogeneity (ReHo) from rs-fMRI. Raw DICOMS were processed using *fMRIPrep* version 25.0.0 (See full processing details in Supplemental Materials). Processing steps included spatial normalization, brain tissue segmentation, head motion correction and non-linear transformations. Processing was performed in surface space to generate CIFTI outputs so that ReHo could subsequently be computed in cortical surface space. Output files were visually inspected for correct framewise displacement and b0 mapping. To further process timecourses, including despiking, confound selection, confound regression and bandpass filtering, we ran XCP-D, which computed vertex-wise ReHo via Kendall’s coefficient of concordance using a 2D mesh grid model across the cortical surface, with each vertex having four neighbors ^37,46^. ReHo was then visually inspected before extracting ROIs. The MNI ROI mask was converted to fsaverage surface space, overlaid onto each CIFTI file, then cifti_parcellate extracted ROIs for each participant. Since surface-based ReHo can only evaluate cortical regions, we also calculate volumetric ReHo to evaluate subcortical regions. Though surface-based ReHo has been shown to be a more reliable measure of functional organization, volumetric ReHo produces comparable age effects ^37,38^. We re-ran *fMRIPrep* with the same pre-processing steps but with each subject’s T1w space as the input. Due to XCP-D not allowing inputs in T1w space, we manually ran each XCP-D processing step with the same parameters as above following a previous group’s established outline (https://github.com/PennLINC/xcp_d) ^47^. This included despiking, confound regression, FWHM smoothing, nuisance regression, bandpassing, and FD censoring. AFNI’s 3dReho calculated 3D volumetric ReHo with the default neighborhood size of 27 voxels. antsApplyTransforms was used to warp our ROI mask to T1w subject space for each participant, which was obtained from their *fMRIPrep* pre-processed anatomical T1w image. Finally, 3dROIstats was used to extract subcortical ROIs. All scans with framewise displacement greater than 0.3 were excluded from analysis, making the final sample n=147 (10-26yo; 77F).

#### T1w/T2w Data Processing

Intracortical myelin was measured with T1w/T2w ratio obtained from T1 and T2 images. We followed the SPM12 MRTool processing pipeline outlined by prior work (https://github.com/austinboroshok/frontoparietal-plasticity; ^48–50^, which takes BIDS files as input and includes steps of B1+ correction, normalization and ratio calculation. Default template maps were preset to MNI2009a. T1 and T2 images went through intensity non-uniformity bias correction (INU), nonlinear histogram matching external calibration, skull-stripping, and warping to MNI. Output was subsequently warped to MNI2009c in order to align with our ROI mask. ROIs were extracted using 3dROIstats and visually inspected for correct alignment to gray matter. The final sample was n=174 (10-26yo; 92F).

#### ROI Mask Segmentation

We chose a priori ROIs based off previous literature showing consistent and robust activation within frontal, parietal and subcortical regions during the anti-saccade task ^5,51^. The mask was segmented to only gray matter and included eight cortical regions: the dorsal anterior cingulate cortex (dACC), dorsolateral prefrontal cortex (dlPFC), frontal eye field (FEF), inferior frontal gyrus (IFG), pre-supplementary motor area (PreSMA), supplementary eye field (SEF), ventrolateral prefrontal cortex (vlPFC), and posterior parietal cortex (PPC); and two subcortical regions: the caudate and putamen.

### Statistical Analysis

Statistical analyses were conducted using R Statistical Software (R 4.3.3). Age-related changes in behavior and MRI measures were examined using generalized additive models (GAMs) (k=3, fx = false), which allow for the assessment of non-linear age-related trends typically found in our age range ^52^. Statistical models controlled for motion, hemisphere, and a random effect of repeating ID (due to multiple ROIs per model). Model outliers ±2 standard deviations from the mean were excluded. All reported significant p-values survived FDR correction for multiple comparisons, *p* < .05.

We first assessed non-linear age-related improvements in anti-saccade performance. We next assessed non-linear changes in neurite density across development and tested whether neurite density related to anti-saccade performance controlling for non-linear age effects in all ROIs. Considering that neurite density index is strongly correlated with intracortical myelin ^28,29^, we statistically regressed out myelin by taking the residual of a linear model comparing NDI and T1w/T2w ratio in order to isolate the effect of neurite density above the contribution of myelin. The resulting signal is labeled as “residualized neurite density” or “rNDI” in Figures 2 and 4 below (see discussion for further interpretation). We conducted separate analyses with residualized neurite density and myelin only in ROIs that had significant associations with neurite density. Next, we assessed non-linear changes in ReHo across development and associations with anti-saccade performance controlling for non-linear age effects in all ROIs. Additionally, we fit a time-varying effect model (TVEM), used to determine at what time point a non-linear association is significant, to inspect an age by ReHo interaction. Finally, we examined associations between NDI and ReHo controlling for non-linear age effects in all ROIs, followed by their interaction in predicting anti-saccade performance. We conducted model comparison in all ROIs using Akaike Information Criterion (AIC) and analysis of deviance tests. We compared four models (Table 1) that predicted inhibitory control performance, the first having just rNDI as the predictor, the second having just ReHo, the third having both rNDI and ReHo, and the fourth having an interaction of rNDI and ReHo. This approach allowed us to assess whether micro-level variables in isolation or in conjunction best explained variability in anti-saccade performance beyond age. For the regions that showed the fourth, most complex model as being the best fit, we fit a TVEM and examined a moderation of rNDI on ReHo predicting inhibitory control performance, controlling for non-linear age effects.

**Figure 2.**
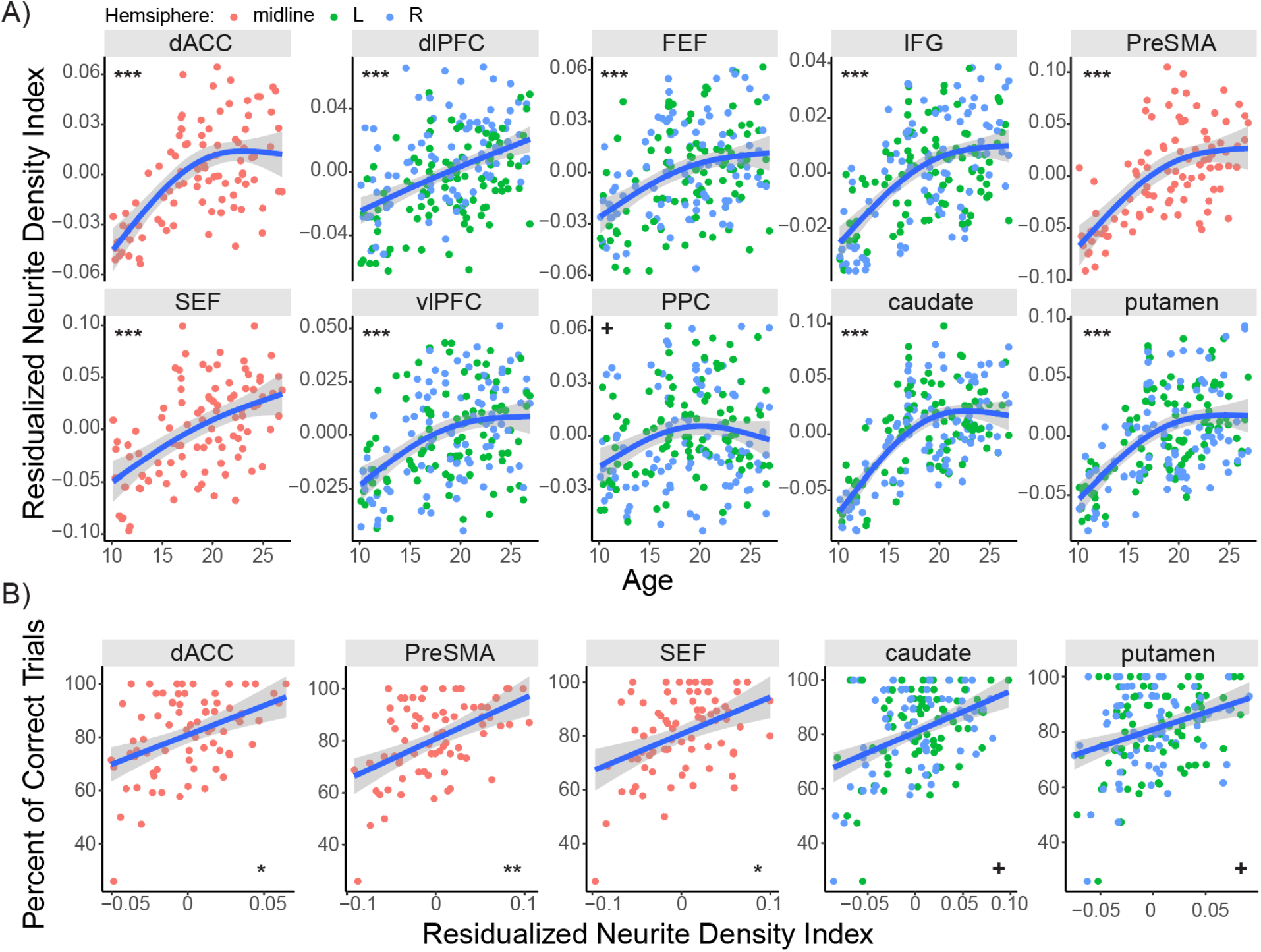
Trajectories of residualized neurite density. Residualized neurite density **A)** increased through development and **B)** was associated with improved inhibitory control accuracy in the dACC, PreSMA, SEF, caudate and putamen. Non-linear age was used as a covariate in addition to controlling for motion, hemisphere, and random effect of repeating ID. Significance survived FDR correction for multiple comparisons. *\*\*p* < .01*, *p* < .05*, +p* < .05 non-residualized NDI.

**Table 1.** Model specifications of best fit for predicting inhibitory control.

| Model | Model specification |
| --- | --- |
| Model 1 | Percent_corr ~ rNDI + s(Age) |
| Model 2 | Percent_corr ~ ReHo + s(Age) |
| Model 3 | Percent_corr ~ rNDI + ReHo + s(Age) |
| Model 4 | Percent_corr ~ rNDI * ReHo + s(Age) |

Note. Percent\_corr, Percent of correct trials on anti-saccade task; ReHo, Regional Homogeneity; rNDI, residualized Neurite Density Index

## Results

### Improvement of inhibitory control through adolescent development

We confirmed age-related improvements in inhibitory control shown in previous literature ^1–7^. We find that performance on the anti-saccade task improves with age, indicated by a higher percent of correct trials (*p* < .001), a decreased latency on correct trials (*p* = .01), and lower trial-to-trial latency variability during correct trials (*p* < .001). Hence, participants show faster, more accurate and more reliable responses as they get older (Figure 1).

### Trajectories of microstructural change related to inhibitory control

We first characterized neurite density across early adolescence through young adulthood. We found NDI significantly increased with age across all cortical and subcortical regions (all *p* < .01). When we residualized for T1w/T2w ratio in our model, all age effects remained significant, with the exception of the PPC (*p* = .1) (Figure 2A; see Figure S1A for non-residualized NDI). Next, we assessed the relationship between microstructure and inhibitory control performance. Greater NDI was associated with improved anti-saccade performance, indicated by an increased proportion of correct responses, in midline-prefrontal regions, including the dACC (*p* = .04), PreSMA (*p* = .03), SEF (*p* = .03), and in the caudate (*p* = .04) and putamen (*p* = .05). This effect persisted after accounting for T1w/T2w ratio (all *p* < .05), with the exception of the putamen (*p* = .16) and caudate (*p* = .08). (Figure 2B; see Figure S1B for non-residualized NDI). See supplementary Figure S1D for confirmatory correlation of T1w/T2w ratio and NDI and Figure S2A for overlaid plots comparing rNDI and NDI. We also found that T1w/T2w ratio increased in our age range across all regions (all *p* < .01), with the exception of the dlPFC (*p* = .12) (Figure S1C). T1w/T2w ratio was not related to anti-saccade performance in any region (all *p* > 0.5).

### Local circuit refinement moderates improved inhibitory control

Next, we investigated how local connectivity matures through adolescence, and how it relates to anti-saccade performance. In alignment with prior evidence, ReHo significantly decreased with age in all cortical and subcortical regions (all *p* < .05), excluding the IFG (*p* = .12) and vlPFC (*p* = .28) (Figure 3A). See supplementary Figure S3A and S3B for parallel evidence of volumetric ReHo decreasing with age and correlating with cortical ReHo. While there were no ROIs showing a direct relationship between local connectivity and inhibitory control, we tested a moderation of ReHo and found that age-related improvement in inhibitory control was moderated by local connectivity in the SEF (*p* = .03), FEF (*p* = .03), and PreSMA (*p* = .03). TVEM demonstrated that within these regions, effects of ReHo on performance were most prominent during early adolescence, such that lower (e.g., more mature) ReHo for one’s age was associated with better inhibitory control (Figure 3B).

**Figure 3.**
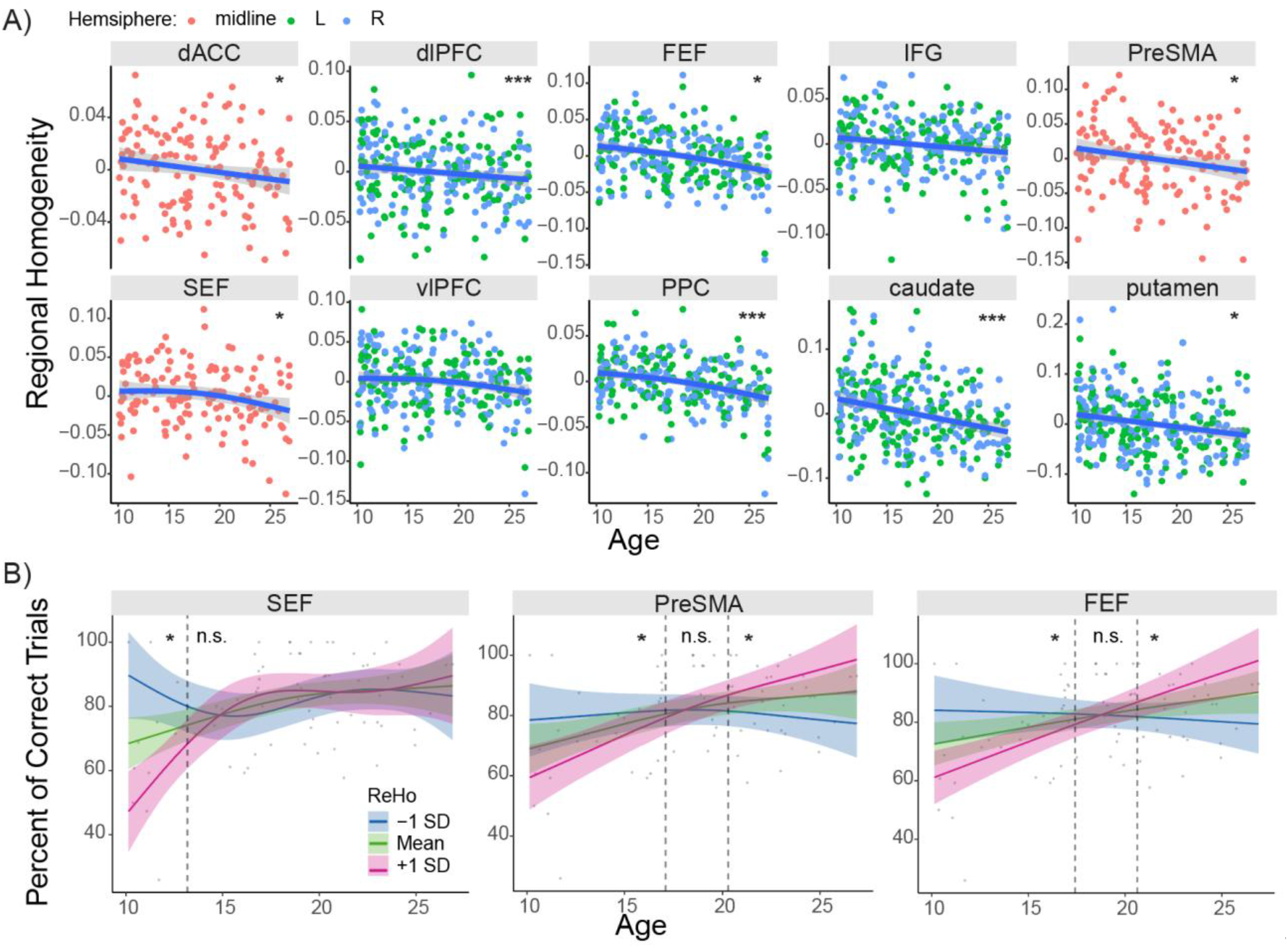
Trajectories of local functional connectivity. **A)** ReHo significantly decreased with age. We controlled for motion, hemisphere, and random effect of repeating ID. **B)** ReHo moderated age-related improvements in inhibitory control accuracy, significant when age was <13yo in the SEF and when age was <17YO and >20yo in the PreSMA and FEF. Significance survived FDR correction for multiple comparisons. \*\*\**p* < .001, \*\**p* < .01, \**p* < .05.

### Interaction of neurite density and local connectivity predict inhibitory control

Finally, we sought to investigate possible associations between developmental changes in microstructure and local circuitry. We found that decreased ReHo was significantly associated with increased NDI, specifically in the FEF (*p* = .003), IFG (*p* = .007), PPC (*p* = .007), caudate (*p* = .003) and putamen (*p* = .01). This effect persisted after accounting for T1w/T2w ratio (all *p* < .01), excluding the putamen (*p* = .07). (Figure 4A; see Figure S4A for non-residualized NDI). Increased T1w/T2w ratio was similarly associated with decreased ReHo, but only in the FEF (*p* = .03) (Figure S4B).

**Figure 4.**
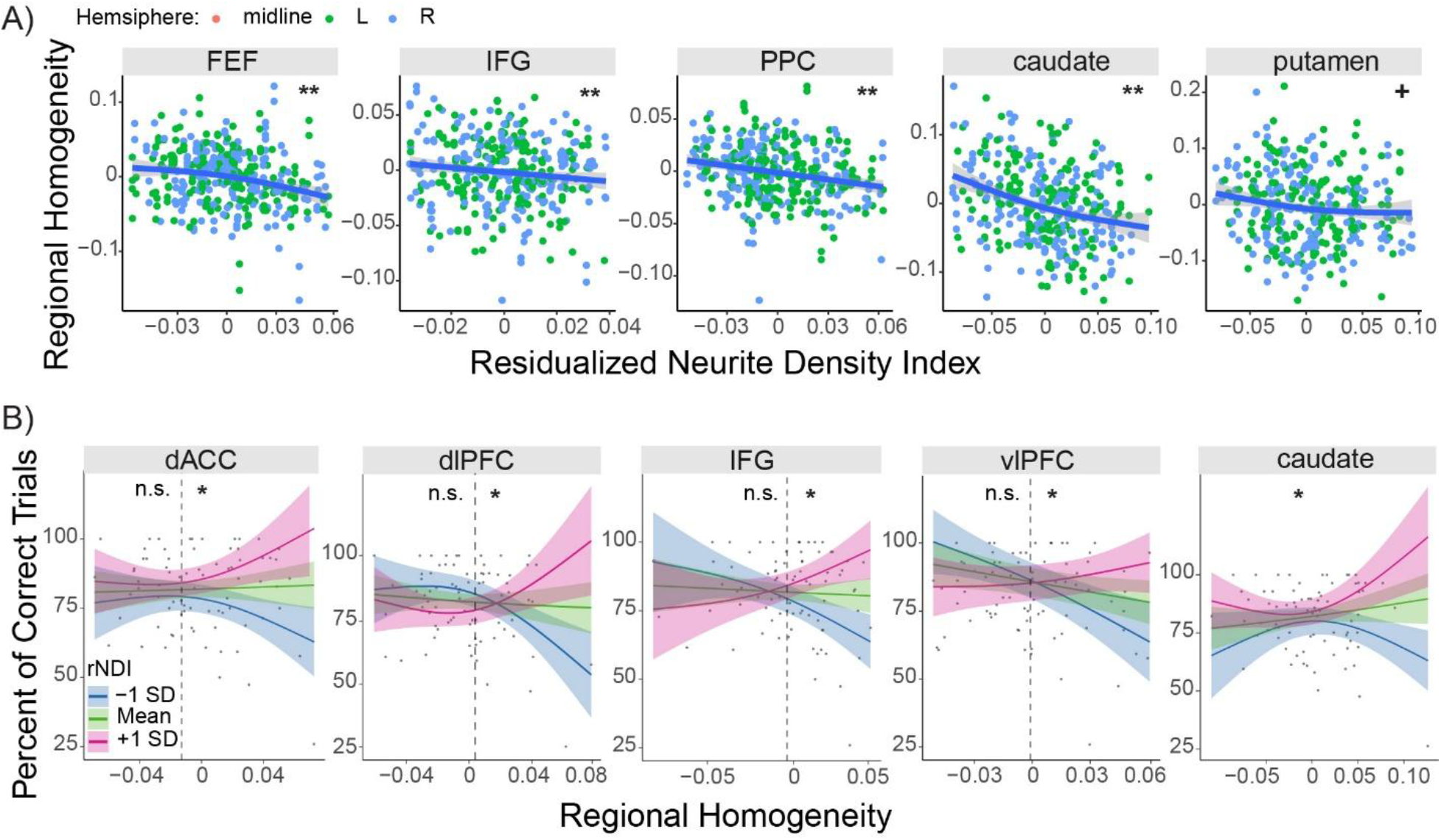
Interaction of neurite density and local connectivity. **A)** Greater residualized neurite density was associated with lower ReHo. Non-linear age was used as a covariate in addition to controlling for motion, hemisphere, and random effect of repeating ID. **B)** There was a moderation of residualized neurite density between local connectivity and inhibitory control accuracy, significant when ReHo > −.01 in the dACC, dlPFC, IFG, vlPFC, and caudate, such that rNDI had the greatest effect on inhibitory control performance when ReHo was high. Non-linear age was used as a covariate. Significance survived FDR correction for multiple comparisons. \*\*\**p* < .001, \*\**p* < .01, \**p* < .05, +*p* < .05 non-residualized NDI.

Further, we assessed how local connectivity and neurite density may interact to support the maturation of inhibitory control. AIC model comparison (see Table 1) revealed that across age in cortical and subcortical regions, an interaction between local connectivity and residualized neurite density predicted developmental improvements in inhibitory control better than either variable alone (Table S1). Based on this result, we tested a moderation of residualized neurite density between local connectivity and inhibitory control and found a significant moderation in the dACC (*p* = .04), dlPFC (*p* = .04), IFG (*p* = .02), vlPFC (*p* = .03), and caudate (*p* = .02). Inspection of a TVEM indicated that within these regions, neurite density had the greatest effect on inhibitory control performance when local connectivity was high (e.g., immature) (Figure 4B).

## Discussion

The present study sought to characterize both the microstructural and functional mechanisms underlying the development of mature inhibitory control through adolescence. First, we aimed to assess developmental trajectories of neurite density and local functional connectivity within gray matter regions and to determine how they each contribute to inhibitory control performance. As hypothesized, NDI, both with and without accounting for myelin content, increased across adolescence, while ReHo showed consistent decreases, in agreement with prior studies ^21–23,37,39^. Further, we found that greater NDI for one’s age and lower ReHo (specifically for young adolescents) was associated with more accurate anti-saccade performance. Lastly, we aimed to understand how microstructure and functional properties interacted to support inhibitory control. We found that rNDI was significantly associated with ReHo independent of age, and the relationship between ReHo and inhibitory control was moderated by rNDI. Our results show that microstructure and local functional circuitry mature in a coordinated manner to support mature cognitive functioning.

We support and expand on preliminary evidence that NDI increases through development. Prior studies have looked at NDI in infancy through adolescence (0-18 yo) ^21–23^ and in specific clinical populations (e.g., schizophrenia, Autism Spectrum Disorder) ^53^, but have not yet characterized gray matter NDI in a broad developmental sample from early adolescence into young adulthood. We investigated gray matter microstructure as an *in vivo* proxy for known microstructural maturation occurring alongside synaptic reorganization, which has been consistently demonstrated in animal and post-mortem work ^13,15^. In addition, only a few studies have shown a direct relationship between gray matter NDI and cognitive functioning ^27,40,54^, and we extend this literature by showing a relationship between NDI and the development of inhibitory control. Most importantly, we interrogate the contribution of myelin in NDI, allowing us to further parse out microstructural properties captured by the NDI signal. It is currently thought that NDI is largely driven by myelin, since it is composed of signal from myelinated axons ^22,28^. Given well-established increases in myelin through development ^30,31,33^, we hypothesized that myelin contributions in the NDI signal were masking other microstructural properties. To address this confound, we statistically controlled for myelin by taking the residual of a linear model with NDI and T1w/T2w ratio and using the residual in our subsequent analyses. These models showed that myelin explained around 20% of variance in NDI, supporting the notion that myelin is partially influencing the calculation of NDI. Importantly, age effects in NDI and associations with ReHo and behavior persisted when using the residualized NDI variable, indicating that age-related myelin contributions alone are not supporting the development of inhibitory control and the remaining microstructural properties are still driving a developmental increase.

Considering what other maturational processes could be driving changes in the NDI signal during the adolescent to adulthood transition, prior literature posits that developmental increases in NDI may reflect axonal growth, greater axonal packing, or greater axonal density ^22,28,55^. Another microstructural element NDI may be capturing is microglial density. A substantial body of literature demonstrates that microglia, resident immune cells in the brain, are critical for synaptic pruning ^56,57^ and that microglial density changes through adolescent development ^58^. Within the NODDI compartment model, however, glia are not present in intra-neurite space, from which NDI is derived, but rather in the extra-neurite space. Another NODDI measure, orientation dispersion index, is derived from both the intra-neurite and extra-neurite space and has been found to positively correlate with histological microglial density ^59^. Still, there is evidence that NDI may be sensitive to changes in microglial density, suggesting that NDI may be informative of not just axonal integrity but microglia-mediated pruning ^60^. For this reason, we hypothesize that NDI may additionally reflect microglial density, among other microstructural properties shaped by synaptic pruning. Future studies are needed to interrogate this hypothesis further by exploring the relationships between our measure of residualized NDI and other *in vivo* proxy measures of glia or synaptic pruning.

Further, our results support previous literature showing that ReHo decreases across adolescence and that it is related to cognitive function ^37,39^. In line with prior evidence showing that a global reduction in ReHo supports the stabilization of working memory across development ^37^, we likewise observed that reductions in ReHo in the FEF, SEF and PreSMA, executive regions of oculomotor control, support more accurate inhibitory control, suggesting a domain-general effect of ReHo on executive function ^7^. Additionally, we found that these reductions in ReHo associate with inhibitory control specifically for young adolescents, suggesting that region specialization, the process of eliminating redundant or inefficient connections within a region through synaptic pruning ^13,38^ may optimize information processing that supports the development of executive function in early adolescence. This is further supported by literature showing developmental decreases in ReHo associating with enhanced intrinsic coding dimensionality, meaning a greater capacity to encode distinct signals within a region ^37^, which is similar to developmental non-human primate studies indicating increases in temporal encoding dimensionality in the prefrontal cortex through adolescence ^61^. Hence, with local neuronal signaling becoming more functionally independent, information processing is optimized to support mature executive functioning for adulthood.

Moreover, we have demonstrated that microstructural elements affect local circuit refinement, as seen by a significant relationship between rNDI and ReHo in both cortical and subcortical regions independent of age. We also found that the relationship between ReHo and inhibitory control was moderated by residualized neurite density, such that the maturity of neurite density had the greatest effect when local connectivity was immature. This underscores a dependence of maturational timing between structure and function during development which together support mature cognitive function. As such, this study is the first to our knowledge to characterize a relationship between local functional connectivity, microstructure, and inhibitory control, providing initial evidence that microstructural elements mature in coordination with local circuitry, contributing to cognitive development. More specifically, our findings show significant associations with more accurate inhibitory control rather than faster inhibitory control performance. Prior work from our lab and others show that latency in the anti-saccade task is reliably decreasing across development and is significantly associated with inter-regional functional connectivity ^51,62^. However, we did not find any relation to latency or latency variability in the present study. This suggests that micro-level mechanisms predominantly support localized computations that execute a goal-driven response, whereas the mechanisms underlying faster processing speed may rely more on the integration of widely distributed, myelinated circuitry.

Across all of the ROIs implicated in inhibitory control, we found that the caudate and putamen most consistently played a significant role in our results. While both are part of the basal ganglia, the caudate receives direct input from frontal cortical oculomotor regions, including the dACC, dlPFC, SEF and FEF, as part of the “associative” loop, whereas the putamen receives input from sensorimotor regions, such as the PreSMA, as part of the “motor” loop ^9,63,64^. The caudate has been implicated in coordinating and planning attentional aspects of behavior, including goal-directed saccades, while the putamen has been implicated in the execution of this planned behavior, though evidence suggests that the putamen also shares responsibility for monitoring eye movement and error ^65^. In the current study, we demonstrate that the caudate and putamen, similar to cortical regions, show significant microstructural and functional maturation. The parallel maturation of cortical and subcortical localized structure and function implies collaborative communication that becomes optimized with development. Furthermore, this may contribute to the efficacy of regions in cortico-striatal loops to functionally specialize, allowing for each region to independently contribute to executing more accurate inhibitory control ^3^. Hence, the maturation of this neural circuitry, by means of region specialization and structural organization, may be central to supporting the maturation of inhibitory control and executive function in general ^66^.

### Limitations

One notable limitation in our study is that the number of subjects included in each analysis was not consistent. Despite our large sample size, approximately 55% (97 participants) had useable data across all four imaging modalities that passed all quality checks. This is due to attrition between behavioral and MRI visits, incompletion of certain scans due to participant discomfort, or excessive movement or other artifacts during MR scanning, which may affect one or more scans. Nevertheless, leveraging multi-modal data with these participants afforded us with an assessment of coordinated microstructural and functional maturation and offered a novel perspective on microstructural mechanisms underlying neurocognitive development. Though current NODDI research has not reached a clear interpretation of NDI, here we disentangle the contribution of intracortical myelin, providing greater specificity to the NDI measurement and a novel hypothesis of NDI reflecting microglial density. In the future, as this was a cross-sectional study, longitudinal data should be leveraged to look at individual differences in development.

## Conclusions

In conclusion, our results suggest that the coordinated maturation of gray matter microstructure and local functional connectivity support the emergence of stable, adult-level inhibitory control. Further, we find that effects of neurite density reflect microstructural properties beyond myelination alone, suggesting another potential microstructural mechanism, such as microglia, that supports region specialization and improved behavior. These results not only inform mechanisms of normative neurocognitive development but can also inform impaired development in psychological conditions, such as mood disorders, substance use disorders, or psychosis, which predominantly emerge during adolescence ^67^. Developmental differences in NDI ^53^ and local connectivity ^68^, along with limitations in inhibitory control ^69,70^ are all evident across these emerging psychopathologies. Hence, at each step— microstructure, function and behavior— we introduce targets for better informing interventions and treatment. Furthermore, this work advances our understanding of the multiple co-sequential processes that, alongside synaptic pruning, contribute to mature inhibitory control and executive functioning.

## Acknowledgements

VOD was supported by NIMH grant R37MH080243 and the Staunton Farm Foundation, both of which BL is the recipient. **VOD:** Conceptualization, Data Curation, Formal Analysis, Investigation, Methodology, Software, Visualization, Writing — original draft, Writing — review & editing. **VJS:** Data Curation, Methodology, Software, Writing — review & editing. **WF:** Data Curation, Methodology, Software, Writing — review & editing. **FJC:** Conceptualization, Funding Acquisition, Methodology, Software, Writing — review & editing. **BL:** Conceptualization, Funding Acquisition, Supervision, Writing — review & editing. Data collection was expertly performed by Hannah Creely, Angela Martinez, Piya Verma and Alyssa Famalette. Preliminary findings from this project have been presented as a poster twice. We thank personnel at the Magnetic Resonance Research Center (MRRC) at UPMC Presbyterian for their assistance in performing the 3T imaging acquisitions. We thank the University of Pittsburgh Clinical and Translational Science Institute (CTSI) for their support in recruiting participants, as well as their support by NIH grant UL1TR001857. We thank members of the lab including Daniel Petrie and Ashley Parr for their guidance and contribution toward editing this manuscript, and Abigail Beatty and Destiny Wright for their daily support and motivation.

## Institution where work was performed

University of Pittsburgh Medical Center. All study procedures were approved by the institutional review board at University of Pittsburgh Medical Center and carried out in accordance with the Declaration of Helsinki.

## Author statement

All authors have seen and approved this manuscript.

