## Supplemental Materials for "Developmental Changes in Gray Matter Microstructure and Local Brain Connectivity Support Inhibitory Control from Adolescence to Young Adulthood"

Victoria O. Dionisos[1]

Valerie J. Sydnor, PhD[2]

Will Foran[2]

Finnegan J. Calabro, PhD[2,3]

Beatriz Luna, PhD[1,2,3]

1. Department of Psychology, University of Pittsburgh, Pittsburgh, PA
2. Department of Psychiatry, University of Pittsburgh, Pittsburgh, PA
3. Department of Bioengineering, University of Pittsburgh, Pittsburgh, PA

**Correspondence:**
Victoria O. Dionisos
Laboratory of Neurocognitive Development
University of Pittsburgh
121 Meyran Ave
Pittsburgh, PA 15213


### 1. Supplemental Figures

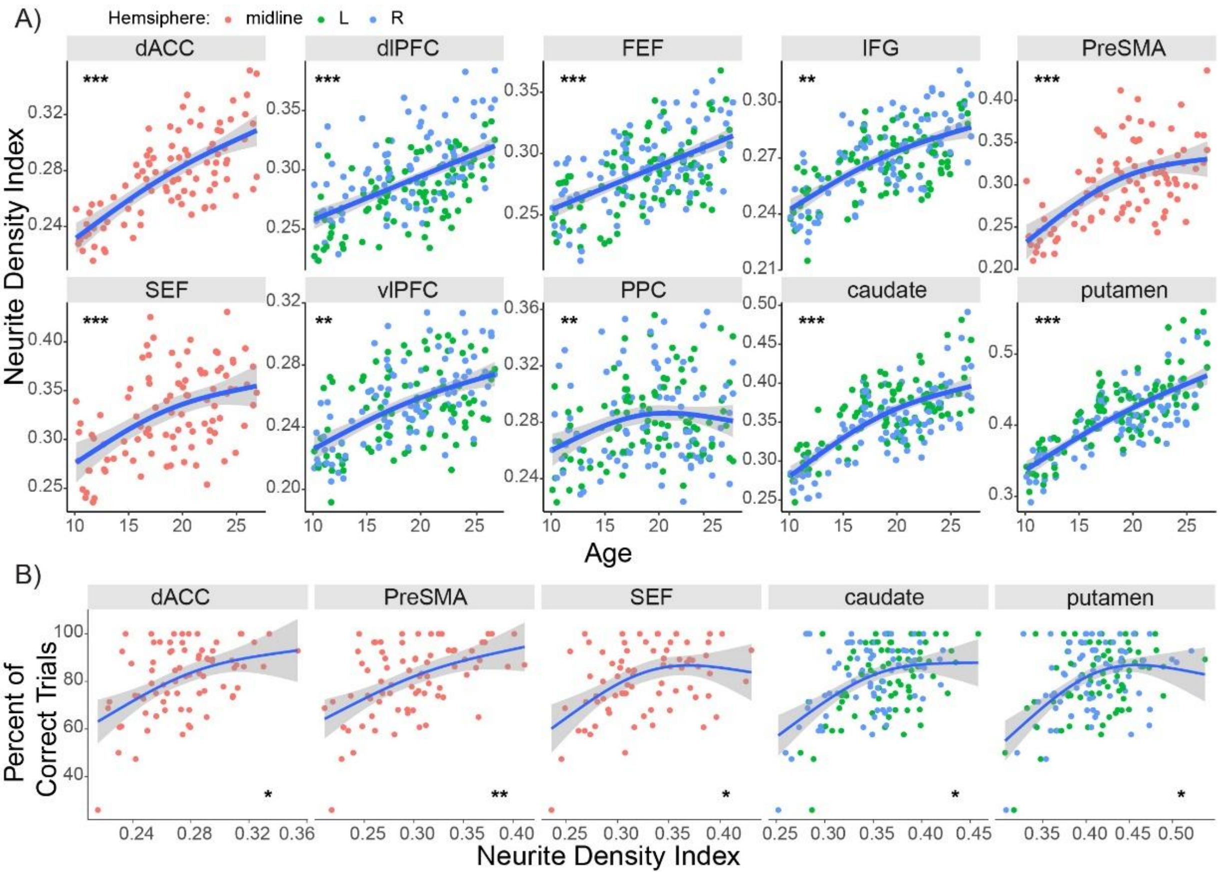

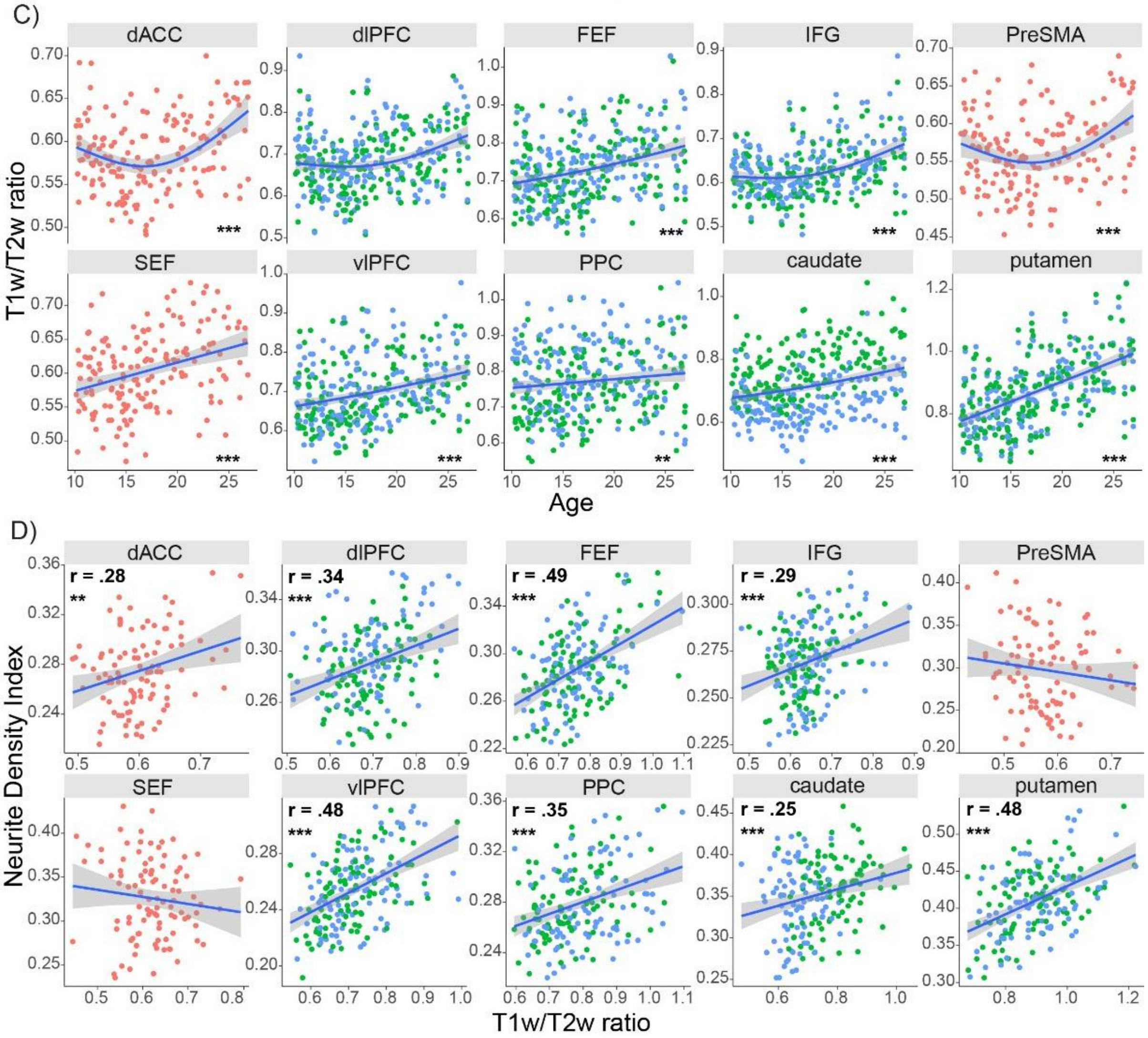


***Figure S1***. Neurite density index **A)** increased through development and **B)** was associated with inhibitory control accuracy. Non-linear age was used as a covariate in addition to controlling for motion, hemisphere, and random effect of repeating ID. As shown in prior studies, **C)** T1/T2w ratio increased with age and **D)** significantly correlated with neurite density index. Significance survived FDR correction for multiple comparisons. $^{***}p < .001$, $^{**}p < .01$

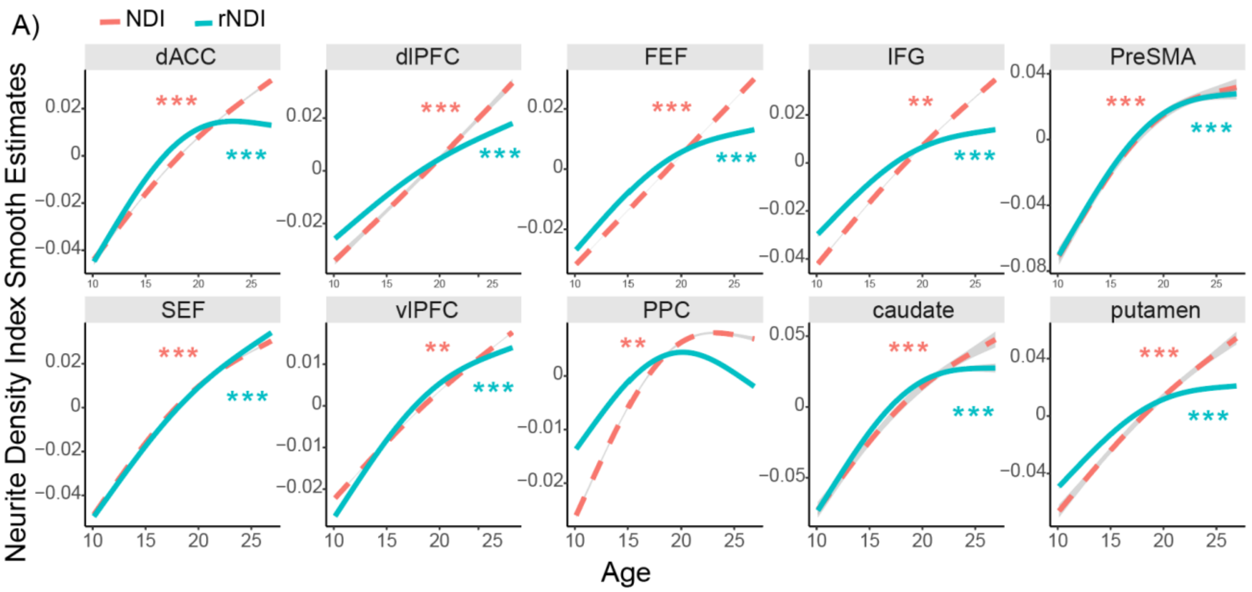


***Figure S2.*** **A)** Smooth estimates of non-residualized NDI and residualized NDI increased through development, with residualized neurite density showing a more pronounced non-linear increase. We controlled for motion, hemisphere, and random effect of repeating ID. Significance survived FDR correction for multiple comparisons. ***$p < .001$, **$p < .01$

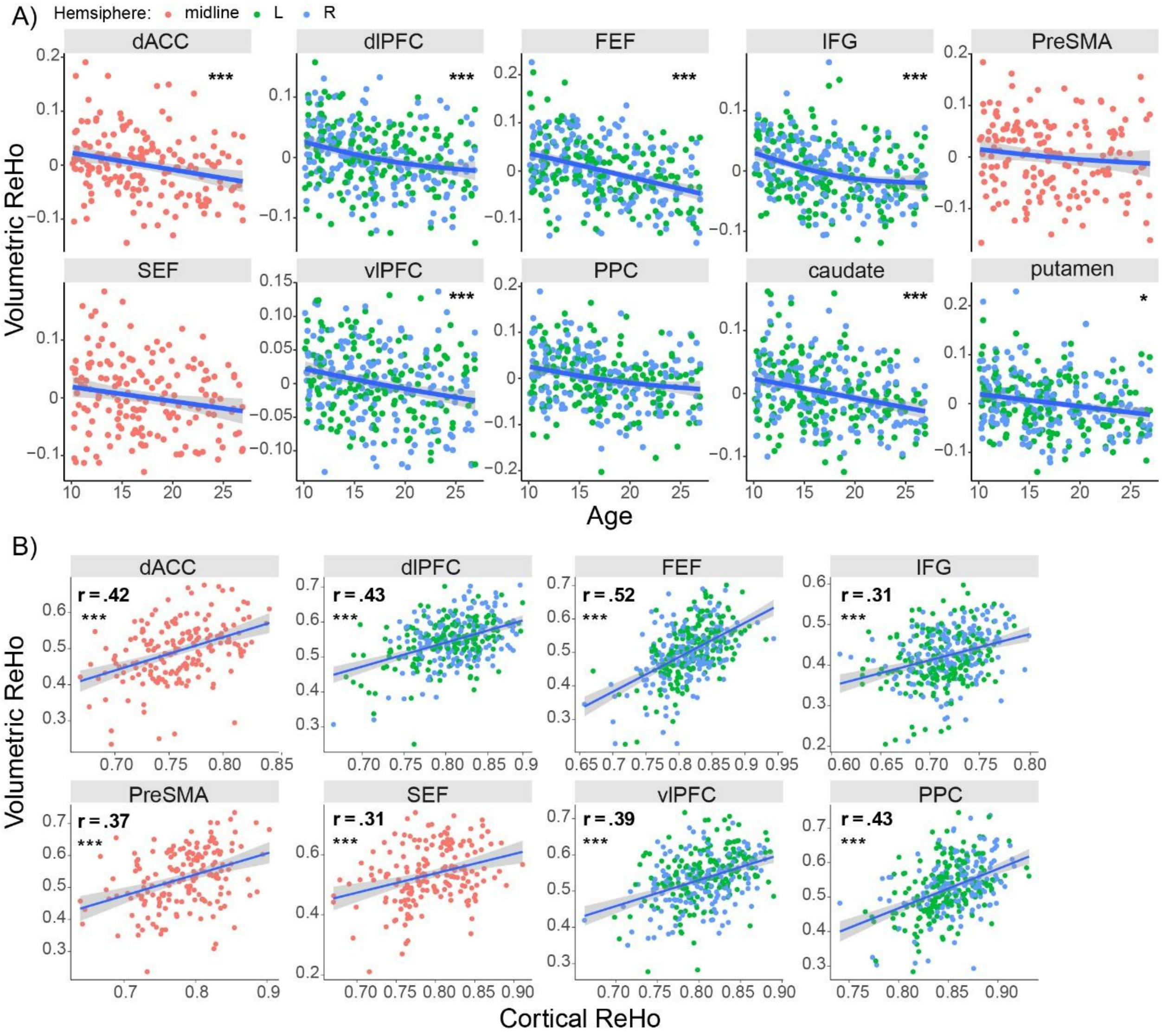


***Figure S3.*** **A)** Volumetric ReHo decreased with age and **B)** significantly correlated with cortical ReHo. We controlled for motion, hemisphere, and random effect of repeating ID. Significance survived FDR correction for multiple comparisons. $***p < .001$, $**p < .01$, $*p < .05$.

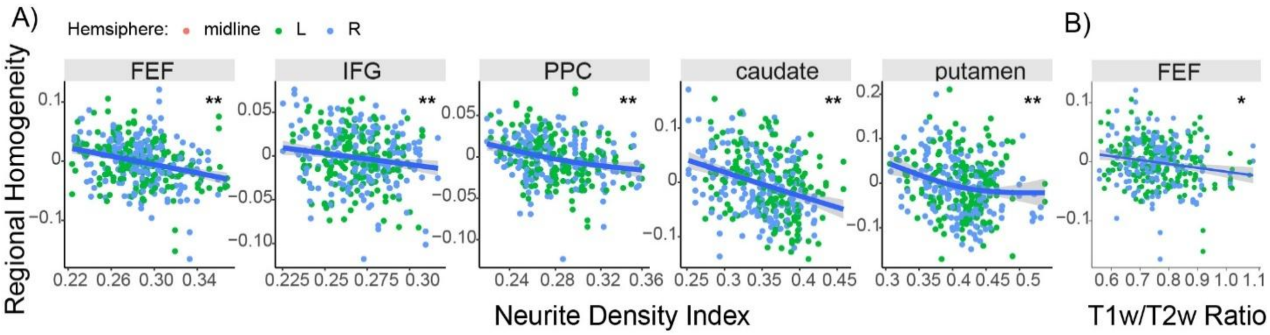


***Figure S4.*** **A)** Neurite density was associated with local connectivity in the FEF, IFG, PPC, caudate and putamen. Non-linear age was used as a covariate in addition to controlling for motion, hemisphere, and random effect of repeating ID. **B)** Increasing T1w/T2w ratio was associated with decreased ReHo in the FEF, with non-linear age as a covariate. Significance survived FDR correction for multiple comparisons. ***$p < .001$, **$p < .01$, *$p < .05$.

### 2. Supplemental Tables

***Table S1.*** AIC model comparison results and analysis of deviance test in regions that show best fit with most complex model. The lowest AIC index is in bold. *p* indicates significance after FDR correction for multiple comparisons. ***$p < .001$, **$p < .01$, *$p < .05$.

| ROI | Model | AIC | Resid DF | Resid Dev | DF | Deviance | *p* |
|---|---|---|---|---|---|---|---|
| dACC | Model 1- NDI | 608 | 69 | 11641 | 3.9 | 1559 | .09 |
| | Model 2- ReHo | 611 | | | | | |
| | Model 3- NDI + ReHo | 610 | | | | | |
| | **Model 4 NDI * ReHo** | **606** | | | | | |
| dlPFC | Model 1- NDI | 618 | 70 | 11426 | 2.8 | 2447 | ** |
| | Model 2- ReHo | 616 | | | | | |
| | Model 3- NDI + ReHo | 618 | | | | | |
| | **Model 4 NDI * ReHo** | **610** | | | | | |
| IFG | Model 1- NDI | 626 | 71 | 11055 | 3.1 | 2681 | ** |
| | Model 2- ReHo | 626 | | | | | |
| | Model 3- NDI + ReHo | 626 | | | | | |
| | **Model 4 NDI * ReHo** | **616** | | | | | |
| SEF | Model 1- NDI | 604 | 68 | 11282 | 3.9 | 1633 | .07 |
| | Model 2- ReHo | 609 | | | | | |
| | Model 3- NDI + ReHo | 605 | | | | | |
| | **Model 4 NDI * ReHo** | **599** | | | | | |
| vlPFC | Model 1- NDI | 619 | 67 | 10637 | 5.2 | 3338 | ** |
| | Model 2- ReHo | 616 | | | | | |
| | Model 3- NDI + ReHo | 618 | | | | | |
| | **Model 4 NDI * ReHo** | **607** | | | | | |
| caudate | Model 1- NDI | 601 | 68 | 11119 | 4.0 | 1959 | * |
| | Model 2- ReHo | 604 | | | | | |
| | Model 3- NDI + ReHo | 603 | | | | | |
| | **Model 4 NDI * ReHo** | **595** | | | | | |

Note. DF, degrees of freedom.

### 3. Supplemental Materials

Results included in this manuscript come from preprocessing performed using *fMRIPrep* 25.0.0 (Esteban et al. (2019); Esteban et al. (2018); RRID:SCR_016216), which is based on *Nipype* 1.9.2 (K. Gorgolewski et al. (2011); K. J. Gorgolewski et al. (2018); RRID:SCR_002502).

#### Preprocessing of B0 inhomogeneity mappings

A total of 5 fieldmaps were found available within the input BIDS structure for this particular subject. A *B0*-nonuniformity map (or *fieldmap*) was estimated based on two (or more) echo-planar imaging (EPI) references with topup (Andersson, Skare, and Ashburner (2003); FSL None).

#### Anatomical data preprocessing

A total of 1 T1-weighted (T1w) images were found within the input BIDS dataset. The T1w image was corrected for intensity non-uniformity (INU) with N4BiasFieldCorrection (Tustison et al. 2010), distributed with ANTs 2.5.4 (Avants et al. 2008, RRID:SCR_004757), and used as T1w-reference throughout the workflow. The T1w-reference was then skull-stripped with a *Nipype* implementation of the antsBrainExtraction.sh workflow (from ANTs), using OASIS30ANTs as target template. Brain tissue segmentation of cerebrospinal fluid (CSF), white-matter (WM) and gray-matter (GM) was performed on the brain-extracted T1w using fast (FSL (version unknown), RRID:SCR_002823, Zhang, Brady, and Smith 2001). Brain surfaces were reconstructed using recon-all (FreeSurfer 7.3.2, RRID:SCR_001847, Dale, Fischl, and Sereno 1999), and the brain mask estimated previously was refined with a custom variation of the method to reconcile ANTs-derived and FreeSurfer-derived segmentations of the cortical gray-matter of Mindboggle (RRID:SCR_002438, Klein et al. 2017). A T2-weighted image was used to improve pial surface refinement. Brain surfaces were reconstructed using recon-all (FreeSurfer

7.3.2, RRID:SCR_001847, Dale, Fischl, and Sereno 1999), and the brain mask estimated previously was refined with a custom variation of the method to reconcile ANTs-derived and FreeSurfer-derived segmentations of the cortical gray-matter of Mindboggle (RRID:SCR_002438, Klein et al. 2017). Volume-based spatial normalization to two standard spaces (MNI152NLin2009cAsym, MNI152NLin6Asym) was performed through nonlinear registration with antsRegistration (ANTs 2.5.4), using brain-extracted versions of both T1w reference and the T1w template. The following templates were were selected for spatial normalization and accessed with *TemplateFlow* (24.2.2, Ciric et al. 2022): *ICBM 152 Nonlinear Asymmetrical template version 2009c* [Fonov et al. (2009), RRID:SCR_008796; TemplateFlow ID: MNI152NLin2009cAsym], *FSL's MNI ICBM 152 non-linear 6th Generation Asymmetric Average Brain Stereotaxic Registration Model* [Evans et al. (2012), RRID:SCR_002823; TemplateFlow ID: MNI152NLin6Asym]. *Grayordinate* "dscalar" files containing 91k samples were resampled onto fsLR using the Connectome Workbench (Glasser et al. 2013).

**Functional data preprocessing**

For each of the 5 BOLD runs found per subject (across all tasks and sessions), the following preprocessing was performed. First, a reference volume was generated, using a custom methodology of *fMRIPrep*, for use in head motion correction. Head-motion parameters with respect to the BOLD reference (transformation matrices, and six corresponding rotation and translation parameters) are estimated before any spatiotemporal filtering using mcflirt (FSL , Jenkinson et al. 2002). The estimated *fieldmap* was then aligned with rigid-registration to the target EPI (echo-planar imaging) reference run. The field coefficients were mapped on to the reference EPI using the transform. The BOLD reference was then co-registered to the T1w reference using bbregister (FreeSurfer) which implements boundary-based registration (Greve and Fischl 2009). Co-registration was configured with six degrees of freedom. The aligned T2w image was used for initial co-registration.Several confounding time-series were calculated based on the *preprocessed BOLD*: framewise displacement (FD), DVARS and three region-wise global signals. FD was computed using two formulations following Power (absolute sum of relative motions, Power et al. (2014)) and Jenkinson (relative root mean square displacement between affines, Jenkinson et al. (2002)). FD and DVARS are calculated for each functional run, both using their implementations in *Nipype* (following the definitions by Power et al. 2014). The three global signals are extracted within the CSF, the WM, and the whole-brain masks. Additionally, a set of physiological regressors were extracted to allow for component-based noise correction (*CompCor*, Behzadi et al. 2007). Principal components are estimated after high-pass filtering the *preprocessed BOLD* time-series (using a discrete cosine filter with 128s cut-off) for the two *CompCor* variants: temporal (tCompCor) and anatomical (aCompCor). tCompCor components are then calculated from the top 2% variable voxels within the brain mask. For aCompCor, three probabilistic masks (CSF, WM and combined CSF+WM) are generated in anatomical space. The implementation differs from that of Behzadi et al. in that instead of eroding the masks by 2 pixels on BOLD space, a mask of pixels that likely contain a volume fraction of GM is subtracted from the aCompCor masks. This mask is obtained by dilating a GM mask extracted from the FreeSurfer's *aseg* segmentation, and it ensures components are not extracted from voxels containing a minimal fraction of GM. Finally, these masks are resampled into BOLD space and binarized by thresholding at 0.99 (as in the original implementation). Components are also calculated separately within the WM and CSF masks. For each CompCor decomposition, the *k* components with the largest singular values are retained, such that the retained components' time series are sufficient to explain 50 percent of variance across the nuisance mask (CSF, WM, combined, or temporal). The remaining components are dropped from consideration. The head-motion estimates calculated in the correction step were also placed within the corresponding confounds file. The confound time series derived from head motion estimates and global signals were expanded with the inclusion of temporal derivatives and quadratic terms for each (Satterthwaite et al. 2013). Frames that exceeded a threshold of 0.5 mm FD or 1.5 standardized DVARS were annotated as motion outliers. Additional nuisance timeseries are calculated by means of principal components analysis of the signal found within a thin band (*crown*) of voxels around the edge of the brain, as proposed by (Patriat, Reynolds, and Birn 2017). The BOLD time-series were resampled onto the left/right-symmetric template “fsLR” using the Connectome Workbench (Glasser et al. 2013). *Grayordinates* files (Glasser et al. 2013) containing 91k samples were also generated with surface data transformed directly to fsLR space and subcortical data transformed to 2 mm resolution MNI152NLin6Asym space. All resamplings can be performed with *a single interpolation step* by composing all the pertinent transformations (i.e. head-motion transform matrices, susceptibility distortion correction when available, and co-registrations to anatomical and output spaces). Gridded (volumetric) resamplings were performed using nitransforms, configured with cubic B-spline interpolation.

**Functional data preprocessing**

For each of the 5 BOLD runs found per subject (across all tasks and sessions), the following preprocessing was performed. First, a reference volume was generated from the shortest echo of the BOLD run, using a custom methodology of *fMRIPrep*, for use in head motion correction. Head-motion parameters with respect to the BOLD reference (transformation matrices, and six corresponding rotation and translation parameters) are estimated before any spatiotemporal filtering using mcflirt (FSL , Jenkinson et al. 2002). The estimated *fieldmap* was then aligned with rigid-registration to the target EPI (echo-planar imaging) reference run. The field coefficients were mapped on to the reference EPI using the transform. The BOLD reference was then co-registered to the T1w reference using bbregister (FreeSurfer) which implements boundary-based registration (Greve and Fischl 2009). Co-registration was configured with six degrees of freedom. The aligned T2w image was used for initial co-registration.Several confounding time-series were calculated based on the *preprocessed BOLD*: framewise displacement (FD), DVARS and three region-wise global signals. FD was computed using two formulations following Power (absolute sum of relative motions, Power et al. (2014)) and Jenkinson (relative root mean square displacement between affines, Jenkinson et al. (2002)). FD and DVARS are calculated for each functional run, both using their implementations in *Nipype* (following the definitions by Power et al. 2014). The three global signals are extracted within the CSF, the WM, and the whole-brain masks. Additionally, a set of physiological regressors were extracted to allow for component-based noise correction (*CompCor*, Behzadi et al. 2007). Principal components are estimated after high-pass filtering the *preprocessed BOLD* time-series (using a discrete cosine filter with 128s cut-off) for the two *CompCor* variants: temporal (tCompCor) and anatomical (aCompCor). tCompCor components are then calculated from the top 2% variable voxels within the brain mask. For aCompCor, three probabilistic masks (CSF, WM and combined CSF+WM) are generated in anatomical space. The implementation differs from that of Behzadi et al. in that instead of eroding the masks by 2 pixels on BOLD space, a mask of pixels that likely contain a volume fraction of GM is subtracted from the aCompCor masks. This mask is obtained by dilating a GM mask extracted from the FreeSurfer's *aseg* segmentation, and it ensures components are not extracted from voxels containing a minimal fraction of GM. Finally, these masks are resampled into BOLD space and binarized by thresholding at 0.99 (as in the original implementation). Components are also calculated separately within the WM and CSF masks. For each CompCor decomposition, the *k* components with the largest singular values are retained, such that the retained components' time series are sufficient to explain 50 percent of variance across the nuisance mask (CSF, WM, combined, or temporal). The remaining components are dropped from consideration. The head-motion estimates calculated in the correction step were also placed within the corresponding confounds file. The confound time series derived from head motion estimates and global signals were expanded with the inclusion of temporal derivatives and quadratic terms for each (Satterthwaite et al. 2013). Frames that exceeded a threshold of 0.5 mm FD or 1.5 standardized DVARS were annotated as motion outliers. Additional nuisance timeseries are calculated by means of principal components analysis of the signal found within a thin band (*crown*) of voxels around the edge of the brain, as proposed by (Patriat, Reynolds, and Birn 2017). The BOLD time-series were resampled onto the left/right-symmetric template "fsLR" using the Connectome Workbench (Glasser et al. 2013). *Grayordinates* files (Glasser et al. 2013) containing 91k samples were also generated with surface data transformed directly to fsLR space and subcortical data transformed to 2 mm resolution MNI152NLin6Asym space. All resamplings can be performed with *a single interpolation step* by composing all the pertinent transformations (i.e. head-motion transform matrices, susceptibility distortion correction when available, and co-registrations to anatomical and output spaces). Gridded (volumetric) resamplings were performed using nitransforms, configured with cubic B-spline interpolation.

Many internal operations of *fMRIPrep* use *Nilearn* 0.11.1 (Abraham et al. 2014, RRID:SCR_001362), mostly within the functional processing workflow. For more details of the pipeline, see the section corresponding to workflows in *fMRIPrep*'s documentation.

Copyright Waiver

The above boilerplate text was automatically generated by fMRIPrep with the express intention that users should copy and paste this text into their manuscripts *unchanged*. It is released under the CC0 license.

Dale, Anders M., Bruce Fischl, and Martin I. Sereno. 1999. "Cortical Surface-Based Analysis: I.

Segmentation and Surface Reconstruction." *NeuroImage* 9 (2): 179–94. https://doi.org/10.1006/nimg.1998.0395.

Esteban, Oscar, Ross Blair, Christopher J. Markiewicz, Shoshana L. Berleant, Craig Moodie, Feilong Ma, Ayse Ilkay Isik, et al. 2018. "fMRIPrep 25.0.0." *Software*. https://doi.org/10.5281/zenodo.852659.

Esteban, Oscar, Christopher Markiewicz, Ross W Blair, Craig Moodie, Ayse Ilkay Isik, Asier Erramuzpe Aliaga, James Kent, et al. 2019. "fMRIPrep: A Robust Preprocessing Pipeline for Functional MRI." *Nature Methods* 16: 111–16. https://doi.org/10.1038/s41592-018-0235-4.

Evans, AC, AL Janke, DL Collins, and S Baillet. 2012. "Brain Templates and Atlases." *NeuroImage* 62 (2): 911–22. https://doi.org/10.1016/j.neuroimage.2012.01.024.

Fonov, VS, AC Evans, RC McKinstry, CR Almli, and DL Collins. 2009. "Unbiased Nonlinear Average Age-Appropriate Brain Templates from Birth to Adulthood." *NeuroImage* 47, Supplement 1: S102. https://doi.org/10.1016/S1053-8119(09)70884-5.

Glasser, Matthew F., Stamatios N. Sotiropoulos, J. Anthony Wilson, Timothy S. Coalson, Bruce Fischl, Jesper L. Andersson, Junqian Xu, et al. 2013. "The Minimal Preprocessing Pipelines for the Human Connectome Project." *NeuroImage*, Mapping the connectome, 80: 105–24. https://doi.org/10.1016/j.neuroimage.2013.04.127.

Gorgolewski, K., C. D. Burns, C. Madison, D. Clark, Y. O. Halchenko, M. L. Waskom, and S. Ghosh. 2011. "Nipype: A Flexible, Lightweight and Extensible Neuroimaging Data Processing Framework in Python." *Frontiers in Neuroinformatics* 5: 13. https://doi.org/10.3389/fninf.2011.00013.

Gorgolewski, Krzysztof J., Oscar Esteban, Christopher J. Markiewicz, Erik Ziegler, David Gage Ellis, Michael Philipp Notter, Dorota Jarecka, et al. 2018. "Nipype." *Software*.

https://doi.org/10.5281/zenodo.596855.

Greve, Douglas N, and Bruce Fischl. 2009. "Accurate and Robust Brain Image Alignment Using Boundary-Based Registration." *NeuroImage* 48 (1): 63–72. https://doi.org/10.1016/j.neuroimage.2009.06.060.

Jenkinson, Mark, Peter Bannister, Michael Brady, and Stephen Smith. 2002. "Improved Optimization for the Robust and Accurate Linear Registration and Motion Correction of Brain Images." *NeuroImage* 17 (2): 825–41. https://doi.org/10.1006/nimg.2002.1132.

Klein, Arno, Satrajit S. Ghosh, Forrest S. Bao, Joachim Giard, Yrjö Häme, Eliezer Stavsky, Noah Lee, et al. 2017. "Mindboggling Morphometry of Human Brains." *PLOS Computational Biology* 13 (2): e1005350. https://doi.org/10.1371/journal.pcbi.1005350.

Patriat, Rémi, Richard C. Reynolds, and Rasmus M. Birn. 2017. "An Improved Model of Motion-Related Signal Changes in fMRI." *NeuroImage* 144, Part A (January): 74–82. https://doi.org/10.1016/j.neuroimage.2016.08.051.

Power, Jonathan D., Anish Mitra, Timothy O. Laumann, Abraham Z. Snyder, Bradley L. Schlaggar, and Steven E. Petersen. 2014. "Methods to Detect, Characterize, and Remove Motion Artifact in Resting State fMRI." *NeuroImage* 84 (Supplement C): 320–41. https://doi.org/10.1016/j.neuroimage.2013.08.048.

Satterthwaite, Theodore D., Mark A. Elliott, Raphael T. Gerraty, Kosha Ruparel, James Loughead, Monica E. Calkins, Simon B. Eickhoff, et al. 2013. "An improved framework for confound regression and filtering for control of motion artifact in the preprocessing of resting-state functional connectivity data." *NeuroImage* 64 (1): 240–56. https://doi.org/10.1016/j.neuroimage.2012.08.052.

Tustison, N. J., B. B. Avants, P. A. Cook, Y. Zheng, A. Egan, P. A. Yushkevich, and J. C. Gee.

2010. “N4ITK: Improved N3 Bias Correction.” *IEEE Transactions on Medical Imaging* 29 (6): 1310–20. https://doi.org/10.1109/TMI.2010.2046908.

Zhang, Y., M. Brady, and S. Smith. 2001. “Segmentation of Brain MR Images Through a Hidden Markov Random Field Model and the Expectation-Maximization Algorithm.” *IEEE Transactions on Medical Imaging* 20 (1): 45–57. https://doi.org/10.1109/42.906424.
